# Pupil dilation indexes – but does not causally influence – conscious error detection: a double-blind, placebo-controlled investigation of performance-monitoring using atomoxetine

**DOI:** 10.1101/2025.11.07.687236

**Authors:** Yoojeong Choo, Adrianna E. Segal, Ryan M. Carnahan, Nathan Chalkley, Ryan Potter, Kylie Dolan, Thomas Nickl-Jockschat, Jan R. Wessel

## Abstract

**Background:** Conscious error detection is accompanied by error-related changes in phasic autonomic activity. This autonomic response is diminished in older age – accompanied by impairments in the conscious detection of action errors – i.e., increased ‘error blindness’. Indeed, the degree to which the autonomic response to errors declines across the lifespan is correlated with the increase in error blindness. However, the direction of causality – whether changes in autonomic reactivity are a consequence or cause of increased error blindness – is still debated. In the present study, we experimentally modulated the phasic autonomic response to action errors in healthy older adults while measuring their conscious error detection.

**Methods:** Across two sessions, thirty healthy older adults (60-80 years old) were given the sNRI atomoxetine or placebo in a double-blind fashion. In each session, they performed an anti-saccade task, which is commonly used to test conscious error detection. The autonomic response to errors was measured via changes in pupil dilation. A novelty-oddball task was also employed as a manipulation check.

**Results:** Atomoxetine reduced phasic pupil dilation to both novel stimuli in the novelty-oddball task and to action errors in the anti-saccade task. However, despite this blunting of the phasic autonomic response to errors, there were no significant differences in conscious error awareness between atomoxetine and placebo. Primary task performance was also unaffected.

**Conclusions:** Despite its effects on phasic autonomic activity after errors, atomoxetine had no effect on conscious error detection in healthy older adults. This suggests that phasic autonomic activity is a consequence, rather than a contributing factor, to conscious error awareness. It also suggests that changes to phasic autonomic activity is unlikely to explain increased error blindness in older age.

## Introduction

The conscious detection of committed action errors is paramount for adaptive behavior and organized goal-directed actions (Mayr, 2004; Wessel, 2012). Conscious error detection is the result of an active decision-making process, in which the brain’s performance-monitoring system gathers multimodal evidence for an error and evaluates this evidence against a set threshold (Steinhauser & Yeung, 2010; Ullsperger et al., 2010). If the threshold is exceeded, widespread neural networks are recruited, resulting in conscious error detection. According to this work, various input signals – neural, behavioral, and autonomic – converge to produce a composite error signal.

The role of autonomic processes in conscious error detection in particular is still debated. Considerable research has shown that consciously perceived errors are accompanied by phasic changes in autonomic activity and that consciously detected errors are accompanied by phasic autonomic arousal (Wessel et al., 2011; Ullsperger et al., 2010; Shalgi et al., 2009; Hajcak et al., 2003). In line with this, the anterior insula cortex (AIC), a key part of the brain’s autonomic network (Critchley et al., 2005; Ferraro et al., 2022), has been suggested as a key node in error awareness and monitoring. According to this and similar work, errors represent “salient” events, which elicit an orienting response (OR) that encompasses the AIC and activates a broader network of brain regions – commonly referred to as the “salience network” (Dosenbach et al., 2006; Seeley et al., 2007; Ullsperger et al., 2010; Wessel, 2018; Menon & Uddin, 2010). The OR is a reflexive cascade of central and autonomic nervous system processes to unexpected environmental changes (Sokolov, 1990). Conscious error awareness and the error-related OR appear to be tightly related, but the direction of causality has yet to be determined – in other words, does a consciously detected error trigger a stronger OR, or does a stronger OR trigger conscious error detection?

Of particular importance to the OR is the locus-coeruleus-norepinephrine (LC-NE) system. Salient stimuli, such as errors, evoke high-frequency phasic activity in the LC, triggering the release of NE, which is crucial for the autonomic response (Morilak et al., 2005). LC-NE activity is closely correlated with task performance (Usher et al., 1998). Pupil diameter is believed to be a sensitive physiological marker of the LC-NE response (Joshi et al., 2016), in line with an increase of pupil diameter found after action errors (Murphy et al., 2011). Indeed, just as other autonomic indices (see above), pupil dilation is increased for consciously perceived action errors compared to errors that went unperceived (Wessel et al., 2011; Critchley et al., 2005; Wessel et al., 2018).

Notably, conscious error detection markedly worsens over the lifespan. Healthy older adults show reduced error awareness even when their primary task performance remains relatively intact (e.g., Harty et al., 2017). Notably, healthy older adults also show blunted autonomic responses to salient stimuli, including errors (Wessel et al., 2018), potentially linking performance monitoring, autonomic responses, and healthy aging. In our previous work, we directly investigated this relationship (Wessel et al., 2018). Healthy older adults showed decreased phasic autonomic activity after errors as well as impaired conscious error detection. Importantly, the degree of this blunted phasic autonomic response correlated with the observed behavioral impairments in conscious error detection. In other words, healthy older adults with greater impairments in their ability to consciously detect their action errors also exhibited smaller phasic autonomic responses, as indexed by pupil dilation.

However, the causal chain of events remains unclear. There are two potential explanations. First, the phasic autonomic response could be a result of conscious error detection, reflecting a saliency response to an error being consciously perceived. Second, the phasic autonomic response could instead be the result of preconscious processes and itself contribute to the decision processes that ultimately lead to conscious error detection (Steinhauser & Yeung, 2010; Ullsperger et al., 2010).

In the present study, we directly tested these two possibilities. Healthy older adults performed two sessions of an anti-saccade task (Hallet, 1978), which reliably elicits both consciously detected and undetected errors (Nieuwenhuis et al., 2001; Wessel et al., 2018; Klein et al., 2007; Endrass et al., 2005). To manipulate autonomic activity, these participants received atomoxetine or placebo in a double-blind, placebo-controlled manner, on separate dates. Atomoxetine is a selective norepinephrine (NE) reuptake inhibitor (sNRI), which modulates both tonic autonomic activity (Pfeffer et al., 2021; O’Callaghan et al., 2025) and the phasic autonomic response (Reynaud et al., 2019; Wernicke et al., 2003). A single, 80 mg dose of atomoxetine has been shown to be effective in altering behavior and performance-monitoring. It leads to heightened phasic alertness and increased error signaling, but also impairs inhibitory control when arousal is elevated (Graf et al., 2011). This could indicate that atomoxetine may contribute to improved error awareness. However, other studies have shown conflicting results, indicating no significant increase in error awareness after a single 60 mg dose of atomoxetine in healthy younger adults (Hester et al., 2012).

Here, we applied atomoxetine to directly influence the phasic autonomic response to errors in the older adults. We expected atomoxetine to increase baseline pupil diameter, and thus reduce further phasic pupil dilation after errors. By pharmacologically manipulating the phasic autonomic response after errors, we aimed to test the two competing hypotheses: If atomoxetine alters the ability of older adults to consciously detect their own action errors, this would indicate that the phasic autonomic response causally contributes to error awareness. If, however, atomoxetine impacts the phasic autonomic response without changing error awareness, this would suggest that phasic autonomic activity is a consequence, not a cause, of error awareness.

## Methods and Materials

### Participants

All procedures were approved by the University of Iowa ethics committee (IRB No. 201805834) and performed in accordance with the Declaration of Helsinki. Thirty healthy older adults from the Iowa City, IA (USA) community were recruited to participate [age mean, *SD* (65.7, 5.0); 27 right-handed, 3 left-handed; 14 females, 16 males]. Inclusion criteria consisted of normal or corrected-to-normal vision and hearing and an age range from 60 to 80 years old. Exclusion criteria included inability to provide consent, current or past history of neurological/psychiatric disorder, marijuana or tobacco use, heavy/frequent alcohol use, actively undergoing treatment with psychoactive medication (i.e. antidepressants, antipsychotics, anxiolytics, stimulants, depressants, mood stabilizers), undergoing treatment with medication with ocular administration (i.e. eye drops) on day of experiment, glaucoma, Mini-Mental Status Examination (MMSE) score under 24, history of autonomic nervous system or cardiac disorders, ongoing treatment with cardioactive medication (i.e. ACE inhibitors, beta-blockers, alpha-blockers, renin/angiotensin system medication), known hypersensitivity to atomoxetine, resting heart rate greater than 90 bpm on day of experiment, resting systolic blood pressure greater than 150 mmHg or diastolic greater than 95 mmHg, atomoxetine-specific contraindications, and history of urinary retention, seizure disorder, liver disease, or coronary/cerebrovascular disease.

### Study Design & Procedure

This was a randomized, double-blind placebo-controlled crossover trial of a single dose of atomoxetine 80 mg or placebo.

Interested individuals were initially recruited over phone/email and scheduled for in-person screening. At the screening visit, they were first evaluated on their ability to sign an informed consent document. If the individual was deemed alert, able to communicate, and understood the consent, they were asked to sign and date. Participants also filled out their demographic information, were tested for compatibility with the eye tracker, completed the MMSE, and underwent a full evaluation by an attending psychiatrist from University of Iowa Health Care and Hospitals (TN-J). Once the individual was cleared for participation in the study, they were prescribed both the placebo and 80 mg atomoxetine dose, as well as scheduled for the two experimental sessions. The first experimental session was planned for at least two weeks after screening, to ensure ample time for the medication to be delivered. The second experimental session was scheduled at least 6 days after the first session. This duration was chosen as the half-life of atomoxetine is about 5 hours in normal metabolizers (and about 24 hours in poor CYP2D6 metabolizers). Thus, 6 days should still be adequate even in poor metabolizers.

Prescriptions were sent to a local compounding pharmacy, where the order of atomoxetine/placebo dispensing was randomized for each participant. The medication was then sent to IDS (Investigational Drug Services) in the University of Iowa Hospitals and Clinics, to be picked up and dispersed to the participant on the day of the study. The experimenter and participant were blinded to whether they were receiving placebo or atomoxetine until the study was concluded. The unblinding list remained with the compounding pharmacy until the end of data collection.

Once the participant arrived for their experimental session, their blood pressure and heart rate were checked by a research nurse. If their resting systolic blood pressure exceeded 150 mmHg, diastolic exceeded 95 mmHg, or heart rate was over 90 bpm, the individual was dismissed. Blood pressure, however, was allowed to be collected multiple times to account for potential “white coat syndrome”, given that the initial screening contained a blood pressure screen that yielded a lower value, and provided the blood pressure value collected on the day of the experiment was higher than typically experienced. The research nurse then administered the medication (placebo or atomoxetine), noted the time and blood pressure of the individual, and set a 110-minute timer. During this waiting period, a Passive Cap EEG system was set up, though EEG data will not be reported in this manuscript. The participant was then read the anti-saccade task instructions. At the 120-minute mark, the research nurse checked blood pressure and heart rate for the second time, ensuring the individual was reacting normally to the medication.

Stimuli were presented using a PC running Windows and Psychtoolbox 3 (Brainard, 1997) under MATLAB 2015b. Stimuli were presented on a 100Hz LCD Monitor (BenQ, 53.5 cm horizontal width) at a viewing distance of 100 cm. Eye-movements and pupil dilation were recorded using a video-based SR Research EyeLink 1000 eye tracker at a sampling rate of 1000Hz and located at a distance of 62cm. A chin and forehead rest were used to maintain a consistent viewing distance and to minimize head movement. Participants responded to the stimuli using their eye movements as well as the computer mouse (for the error detection assessment and confidence rating). The ambient light in the room was kept at a constant, low level for all subjects. The eye tracker was calibrated at the beginning of the experiment and recalibrated after each block of the experiment.

Between blocks of the anti-saccade task, the participants were given 1-2 minutes to rest and review their task performance. After completing 6 blocks of the anti-saccade task, the participant then moved on to the novelty-oddball task. At the end of this task, the research nurse collected blood pressure and heart rate for the third and final time. An attending physician also checked in with the participant before they were dismissed. If it was the individual’s second experimental session, they were debriefed (**Figure 1**).

**Figure 1.**
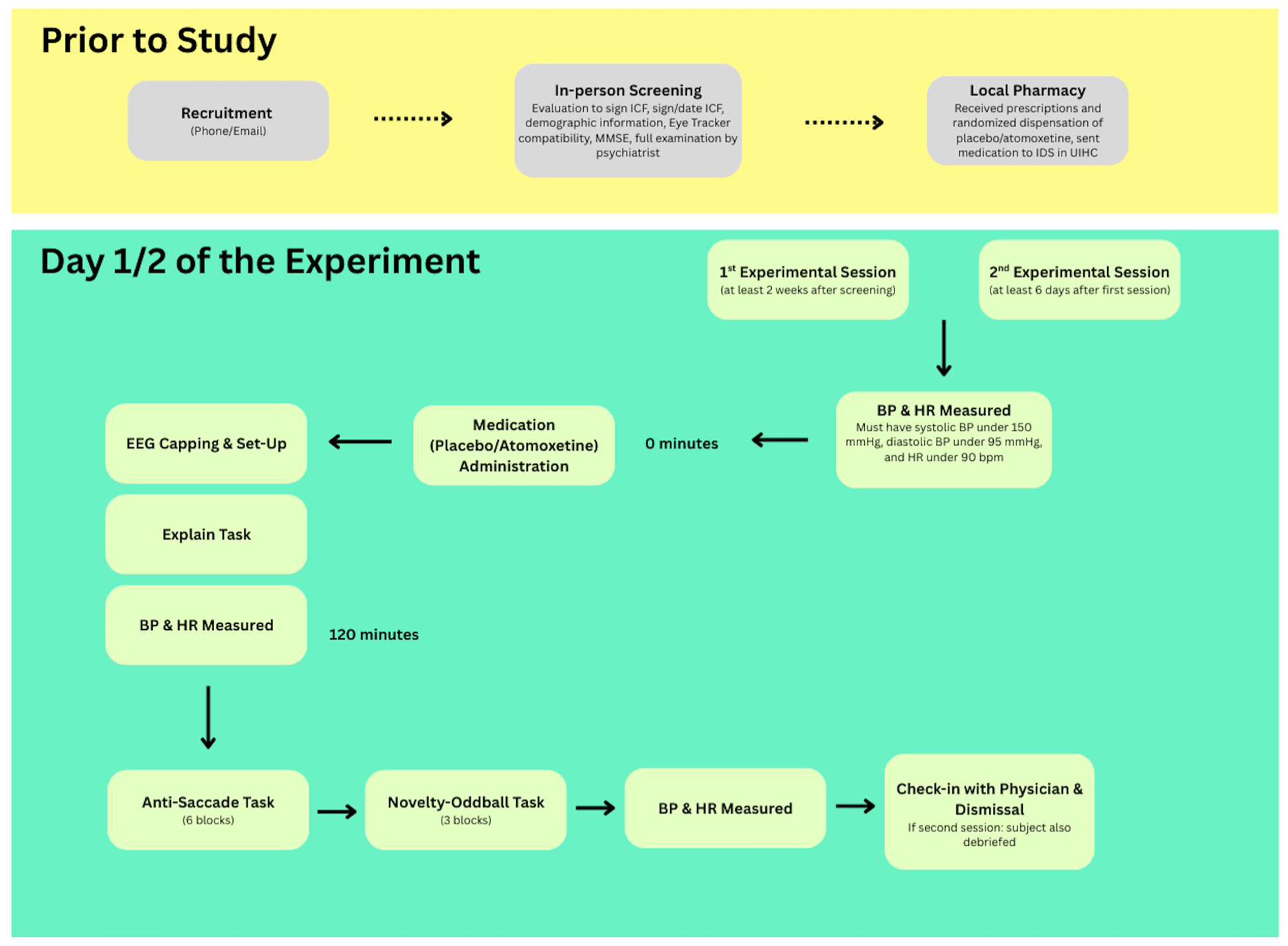
Flowchart of Procedure.

### Main task: Anti-saccade task

The experimental task was designed to induce a reliable amount of undetected action errors. It was furthermore designed to avoid contaminating post-error slowing measurements through the trial-wise error awareness assessment. Participants were instructed to respond to visual stimuli using their eyes and then rate the accuracy of their responses using the computer mouse. The task was identical to that used in Wessel et al., (2018), with one modification. While anti- and prosaccade trials were equally likely in that study, we here used 80% anti-saccade trials, since prosaccade trials produced almost no errors. Some prosaccade trials were retained in the current work to prevent participants from reconfiguring their oculomotor planning towards only producing antisaccades. The remainder of the task is identical to that of Wessel et al. (2018) and the following description is quoted from therein.

An initial fixation cross in the center of the screen (black background) instructed the participants about whether they had to perform a pro- or an anti-saccade on the upcoming trial (green = pro-saccade; red = anti-saccade). The fixation cross was flanked by two yellow boxes (3°x3° of visual angle), presented in the vertical center of the screen, with an offset of +/-10° of visual angle. After 750 ms, the target stimulus, a white disc (diameter: 2°) was displayed in one of the two boxes, and the participants had to respond as quickly as possible by making either a pro-saccade towards the target, or an anti-saccade towards the opposite box. After the 1,000 ms response window was over, a rating screen appeared on a select number of trials. The rating screen appeared on the first correct trial after error commission, after every action error (erroneous pro-saccade in the anti-saccade condition, and vice versa). After every error, a random number between 4 and 10 was generated, and the post-error trial that corresponded to that number (and its subsequent post-trial) was slated for an error awareness query. The automatic algorithm ensured that this trial, as well as the following trial, would be of the same type (anti/pro-saccade) as the trial on which the error was committed and the post-error trial. This achieved two things: first, it ensured an equal number of error detection queries after correct and erroneous responses, and therefore prevented a bias towards error signaling on the part of the participants. Second, it automatically led to matched trial and post-trial types for post-error behavioral comparisons. If the trial that was slated for the error detection query was itself an error, the error detection query was then performed after the next pair of correct trials. The rating screen itself consisted of a simple query (“Error?”) with two response options (“Y”/“N”), one of which the participants had to select by moving the mouse cursor to one of the alternatives and pressing the left mouse button. Participants were instructed that the error detection query pertained to the last-but-one trial (instead of the immediate last trial). That way, reaction times on the post-error and post-correct trials were unaffected by the error detection query. After the initial binary query, a visual analogue scale appeared, which prompted the participants to indicate the subjective certainty of the accuracy assessment by using the mouse to click on the appropriate point of the spectrum that ranged from “unsure” to “very sure”. After the performance rating, participants received feedback about the accuracy of their assessment (“Correct” in green or “Incorrect” in red displayed in the middle of the screen). Participants were incentivized to respond as accurately as possible on the error detection query by receiving a ‘time-out’ on incorrectly rated trials: the feedback on correctly rated trials (hits and reported errors) was on the screen for 1s, whereas the feedback for incorrectly rated trials (false alarms and unreported errors) was on the screen for 5s. After an ITI of 1s (during which only a white fixation cross and the yellow boxes were displayed), the next trial begin with the next cue (red or green fixation cross). Participants performed 600 trials in total, divided across 6 blocks.

**Figure 2.**
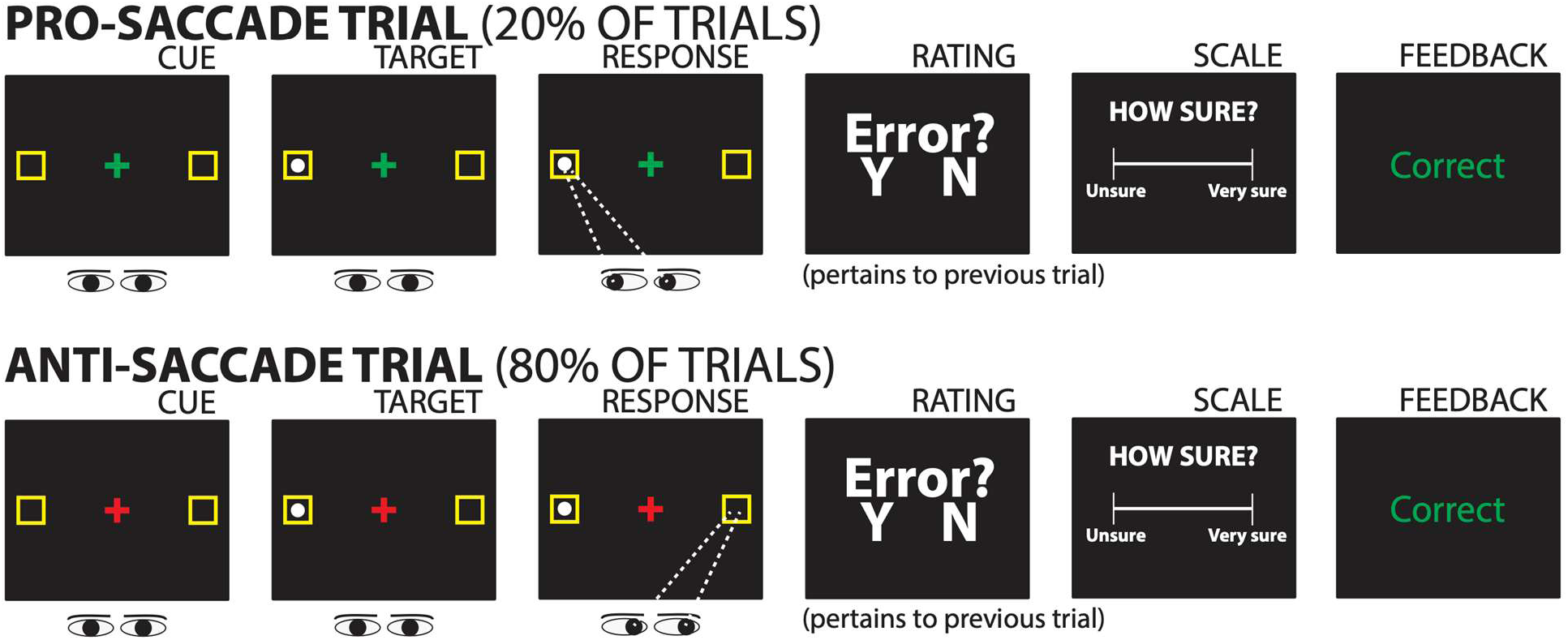
Diagram of the Anti-saccade task.

### Manipulation Check: Novelty-Oddball task

To ensure that the administration of atomoxetine had the predicted effects on autonomic activity, we administered a novelty-oddball task (Courchesne et al., 1975), which is known to reliably evoke a phasic orienting response to unexpected stimuli. Participants were played one of three sounds on each trial: a standard sound (80% of trials), a rare but expected oddball sound (10%) and a rare and unexpected novel sound (10%), which is known to elicit the orienting response. The standard and oddball sounds were 200 ms sine wave tones of two different frequencies (600 and 1,200 Hz, assignment as standard or oddball frequency randomly assigned for each participant). The novel sounds were 200 ms birdsong segments taken from our prior work (Wessel & Aron, 2013). Each trial consisted of a 200 ms sound presented, followed by a 800 ms delay until the next trial. A fixation cross was displayed in the center of the computer screen. Participants were instructed to count the number of oddball sounds in each block of trials. Trials were presented in 3 blocks of 90 trials each (270 total trials, of which 27 were oddball and 27 were novel sounds). Trials were presented in pseudo-random order, with the constraint that each section of 10 trials contained exactly one instance each of a novel and an oddball sound. This was done to ensure that the local probability of novel and oddball sounds remained roughly stable throughout the experiment.

### Pupil data analysis

#### Preprocessing

Pupil data were preprocessed following the pipeline reported in Wessel et al. (2018). The following steps are adapted from that procedure. Data were analyzed using custom scripts in MATLAB 2023b. After conversions into ASCII format using the SR Research’s EDF2ASC tool, raw pupil tracking data were imported into MATLAB and processed in event-related manner for the anti-saccade and novelty-oddball tasks separately. Blinks were interpolated in the continuous trace, except when occurring between stimulus and response in the anti-saccade task, in which case the trial was excluded. In the novelty-oddball task, no responses followed stimulus presentation.

#### Anti-saccade task

Two participants were excluded from the anti-saccade task dataset, one due to a technical defect in one of the two sessions, and one due to chance-level performance, leaving N = 28. Epochs were extracted from -500 to 3500 ms relative to primary response on each trial. Each epoch was baseline-corrected using the 500 ms preceding target onset. To identify the time range during which pupil dilation was modulated by error awareness (Wessel et al., 2011), we conducted a 2 x 3 repeated-measures ANOVA with the factors SESSION and TRIAL TYPE (Hits, Reported errors, Unreported errors) at every sample point from the response until the end of the epoch. This yielded three vectors of p-values (length = 3500), corresponding to the main effects and interaction at each time point. To identify the critical period, we then used the vector for the main effect of TRIAL TYPE and corrected the *p*-values using the false-discovery rate (FDR) procedure (Benjamini et al., 2006) with a threshold of *p* = 0.001. Mean pupil responses were then averaged across this period for each condition and tested using a 2 x 3 repeated-measures ANOVA, followed by planned pairwise comparisons (follow-up *t*-tests). Finally, we tested the association between error awareness rating certainty and pupil dilation by correlating mean certainty ratings for reported errors with pupil responses during those errors using Pearson’s r correlations.

#### Novelty-Oddball task

One participant was excluded due to hardware failure, leaving N = 29. Preprocessing followed the same procedure as described above. Epochs were extracted from -500 to 2500 ms relative to stimulus onset and baseline-corrected using the 500 ms preceding stimulus onset. To determine the time range in which pupil dilation was modulated by stimulus type, we conducted a 2 x 3 repeated-measures ANOVA with the factors SESSION (placebo, atomoxetine) and STIMULUS TYPE (standard, oddball, and novel) at every sample point until the end of the epoch. This yielded three vectors of *p*-values (length = 2500), corresponding to the two main effects and their interaction. The critical period was then defined from the vector for the main effect of STIMULUS TYPE, applying FDR correction (*p* = 0.001; Benjamini et al., 2006). Mean pupil responses were then averaged across this period for each condition and participant and analyzed using a 2 x 3 repeated-measures ANOVA, with planned comparisons conducted using follow-up t-tests. Furthermore, we tested for differences in tonic pupil dilation using the same 2 x 3 repeated-measures ANOVA on the mean pupil dilation during the baseline period. These analyses served as manipulation checks, confirming that both tonic autonomic activity and the phasic autonomic response were significantly altered by atomoxetine.

### Behavioral data analysis

#### Anti-saccade task

Behavioral data were analyzed using our custom scripts in MATLAB 2023b. The analysis pipeline largely followed Wessel et al. (2018), from which the following description is adapted. Trials were excluded if they contained an eyeblink before response emission, a saccadic RT < 80 ms (i.e., anticipatory saccade), or no response (i.e., miss).

Overall task performance was assessed by quantifying error rates and primary saccadic RTs (for both correct and error trials) across all remaining trials. Error detection rates (i.e., detection accuracy), rating certainty, and post-error slowing (PES) were quantified based on trials that included an error detection query after each trial. Error detection rates were computed using signal detection theory (Green & Swets, 1966) as the proportion of correctly detected trials (hits and reported error) relative to all four trial types (hits, false alarms, reported errors, and unreported errors). Error trials were categorized as reported (perceived) and unreported (unperceived) to examine differences between error types. PES was quantified separately for reported and unreported errors in each session as the percent change from correct saccadic RT:

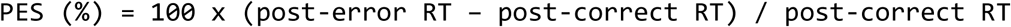

All query-based metrics were expressed as percent change (%) by multiplying the respective ratios by 100.

### Statistical analyses

Error rates, error detection rates, false alarm rates were compared between atomoxetine and placebo sessions using paired-samples *t*-tests (two-sided). Primary saccadic RTs were compared using a 2 x 2 repeated measures ANOVA with the factors SESSION (placebo, atomoxetine) and TRIAL ACCURACY (correct, error) based on all trials. PES was analyzed using a 2 x 2 repeated measures ANOVA with the factors SESSION (placebo, atomoxetine) and ERROR TYPE (reported, unreported). For ANOVAs, post-hoc *t*-tests were performed where indicated and corrected using the Bonferroni-Holm procedure for multiple comparisons. Planned between-session contrasts (placebo vs. atomoxetine) were not corrected for multiple comparisons. Effect sizes are reported as Cohen’s *d* for *t*-tests and partial eta^2^ (*η^2^_p_*) for ANOVAs. When sphericity was violated, Greenhouse-Geisser corrections were applied.

## Results

### Pupil dilation results

#### Atomoxetine modulates the phasic autonomic orienting response in the novelty-oddball task

Because the novelty-oddball task served as a manipulation check of the pharmacological intervention, we first report the pupil dilation results from this task before turning to the main anti-saccade task.

As expected, atomoxetine substantially increased baseline pupil dilation (**Figure 3A**), as revealed by a significant main effect of SESSION (*F*(1,28) = 12.47, *p* = 0.001, *η^2^_p_* = 0.308). There was also an unexpected significant main effect of STIMULUS TYPE on tonic pupil diameter (F(2,56) = 24.36, *p* <.001, *η^2^_p_* = 0.465). This effect was likely due the short ITI used in the novelty oddball task, combined with the fact that each section of 10 trials could only contain one novel and one oddball stimulus each. Therefore, standard events were often preceded by oddball/novel events, which produce pupil dilation (see below), whereas oddball/novel events were rarely preceded by other oddball/novel events. Indeed, removing standard trials that were preceded by oddball/novel events substantially removed the size of the main effect of STIMULUS TYPE. Importantly, there was no interaction between SESSION and STIMULUS TYPE (*F*(2,56) = 0.79, *p* = 0.459, *η^2^_p_* = 0.027).

**Figure 3.**
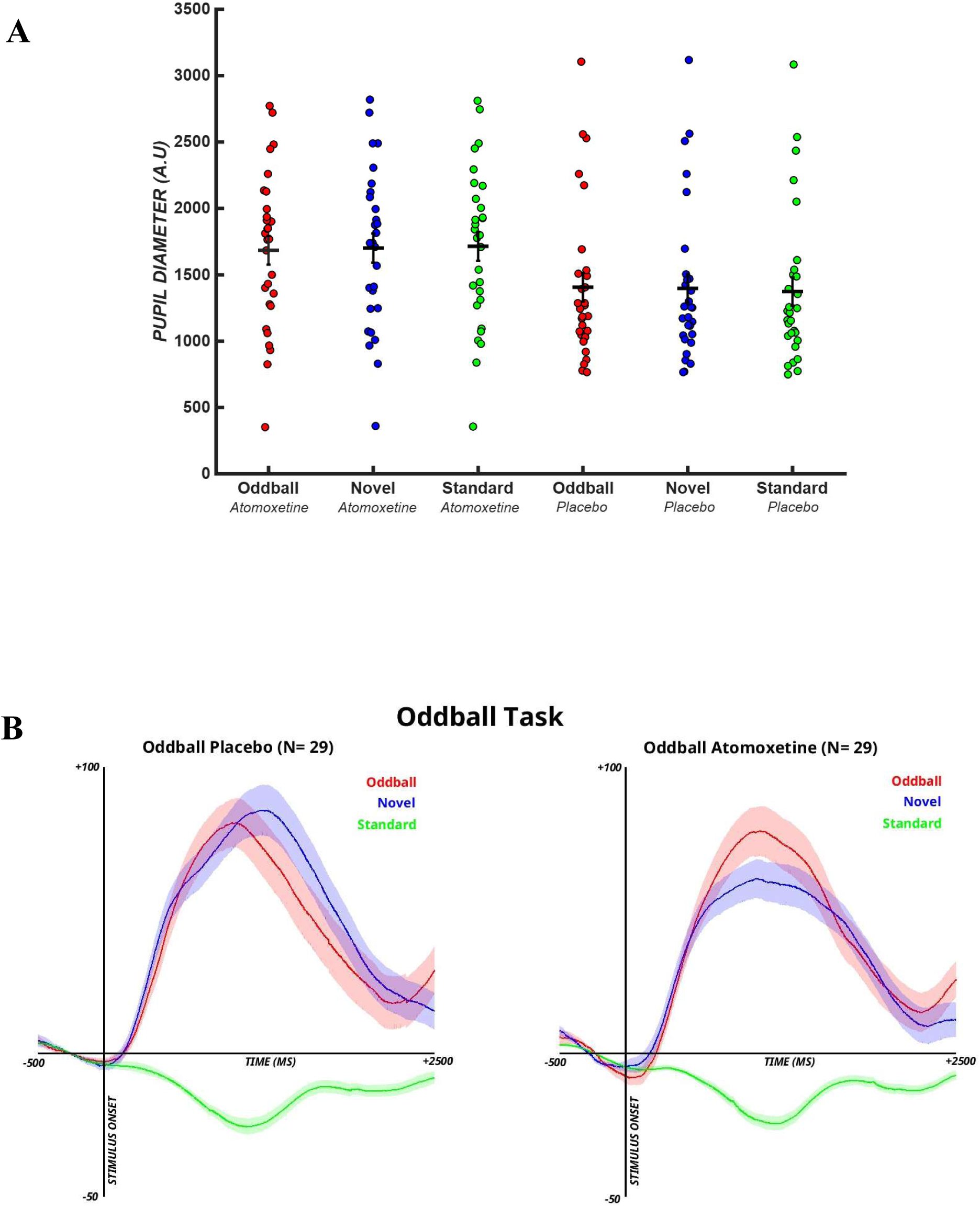
**A) Baseline pupil data during the novelty-oddball task (n = 29)**. On atomoxetine, participants’ baseline pupil diameter was significantly larger than during placebo trials. **B) Grand mean error-related pupil data during the novelty-oddball task (n = 29).** Left: on placebo, subjects displayed an expected increase in pupil diameter after oddball/novel sounds. Right: with atomoxetine administration, the pupil response to novel sounds was reduced.

As expected, atomoxetine also affected the phasic pupil response (**Figure 3B**). A sample-to-sample ANOVA revealed significant effects of STIMULUS TYPE beginning at 191 ms after the stimulus onset and lasting until the end of the epoch (FDR corrected *p* < .001). Based on this analysis, we defined the critical periods as 191-2500 ms, and all subsequent analyses were conducted on the average pupil dilation within this window.

Within this critical period, pupil dilation showed a significant main effect of STIMULUS TYPE (*F*(2, 56) = 59.88, *p* < .001, *η^2^_p_* = 0.584). Post-hoc t-tests (Holm-Bonferroni corrected) confirmed that both oddball (*M* = 44.39) and novel sounds (*M* = 44.49) elicited significantly larger pupil responses than standard tones (*M* = -13.88) (oddball vs. standard; *t*(28) = 8.67, *p* < 0.001, *d* = 2.166; novel vs. standard; *t*(28) = 11.16, *p* < .001, *d* = 2.170), whereas oddball and novel stimuli did not differ (*t*(28) = -0.02, *p* = 0.987, *d* = 0.004). These findings confirm the expected orienting response to rare sounds.

The main effect of SESSION indicated a trend toward reduced pupil responses under atomoxetine (*F*(1, 28) = 3.41, *p* = 0.075, *η^2^_p_* = 0.017). The SESSION x STIMULUS TYPE interaction effect was not significant (*F*(2, 56) = 1.75, *p* = 0.184, *η^2^_p_* = 0.012). Planned comparisons showed that pupil dilation to novel sounds was smaller under atomoxetine (*M* = 38.07, *SD* = 27.07) compared to placebo (*M* = 50.92, *SD* = 33.75) (*t*(28) = -1.74, *p* = 0.046, *d* = 0.323, one-sided test). This pattern is consistent with a reduced phasic autonomic response to surprising stimuli under atomoxetine. This is likely caused by the substantially expanded baseline dilation in the atomoxetine condition (see above). No session differences were found for oddball (atomoxetine: *M* = 43.46, *SD* = 31.88 vs. placebo: *M* = 45.31, *SD* = 36.10; *t*(28) = -0.39, *p* = 0.700, *d* = 0.072) or standard tones (atomoxetine: *M* = -13.28, *SD* = 7.59 vs. placebo: *M* = -14.48, *SD* = 9.60; *t*(28) = 0.65, *p* = 0.521, *d* = 0.121).

#### Atomoxetine also modulates the phasic autonomic response to action errors

Figure 4 shows pupil dilation following responses. As expected, the sample-to-sample ANOVA revealed significant effects of TRIAL TYPE beginning at 1103 ms after the response and lasting until the end of the epoch (FDR corrected *p* < .001). Based on this analysis, we defined the critical period as 1103-3500 ms, and all subsequent analyses were conducted on the average pupil dilation within this window. Within this critical period, a repeated-measures ANOVA revealed a significant main effect of TRIAL TYPE (*F*(2, 54) = 21.80, *p* < 0.001, *η_p_^2^* = 0.447), the main effect of SESSION reached the threshold of significance (*F*(1, 27) = 4.20, *p* = 0.050, *η_p_^2^* = 0.135). The SESSION x TRIAL TYPE interaction was not significant (*F*(2, 54) = 1.31, *p* = 0.278, *η_p_^2^* = 0.046).

Across sessions, post-hoc comparisons (Holm-Bonferroni corrected) replicated the established pattern reported in Wessel et al. (2018). Both reported errors (*M* = 27.24) and unreported errors (*M* = 6.70) elicited significantly greater pupil dilation than correctly identified correct responses, i.e., hits (*M* = -23.42) (reported errors vs. hits; *t*(27) = 6.80, *p* < 0.001, *d* = 0.951; unreported errors vs. hits; *t*(27) = 3.50, *p* = 0.003, *d* = 0.565). Moreover, reported errors produced greater dilation than unreported errors (*t*(27) = 2.93, *p* = 0.007, *d* = 0.385). These results confirm that the expected phasic autonomic response to errors was robustly present in our data. Moreover, we replicated the previously reported associated between certainty ratings for consciously perceived errors and the pupil response to those errors (*r* = 0.41, *p* = 0.03; Figure 5). Subjects with a greater pupil dilation response on reported errors were also more subjectively confident of their error rating.

**Figure 4.**
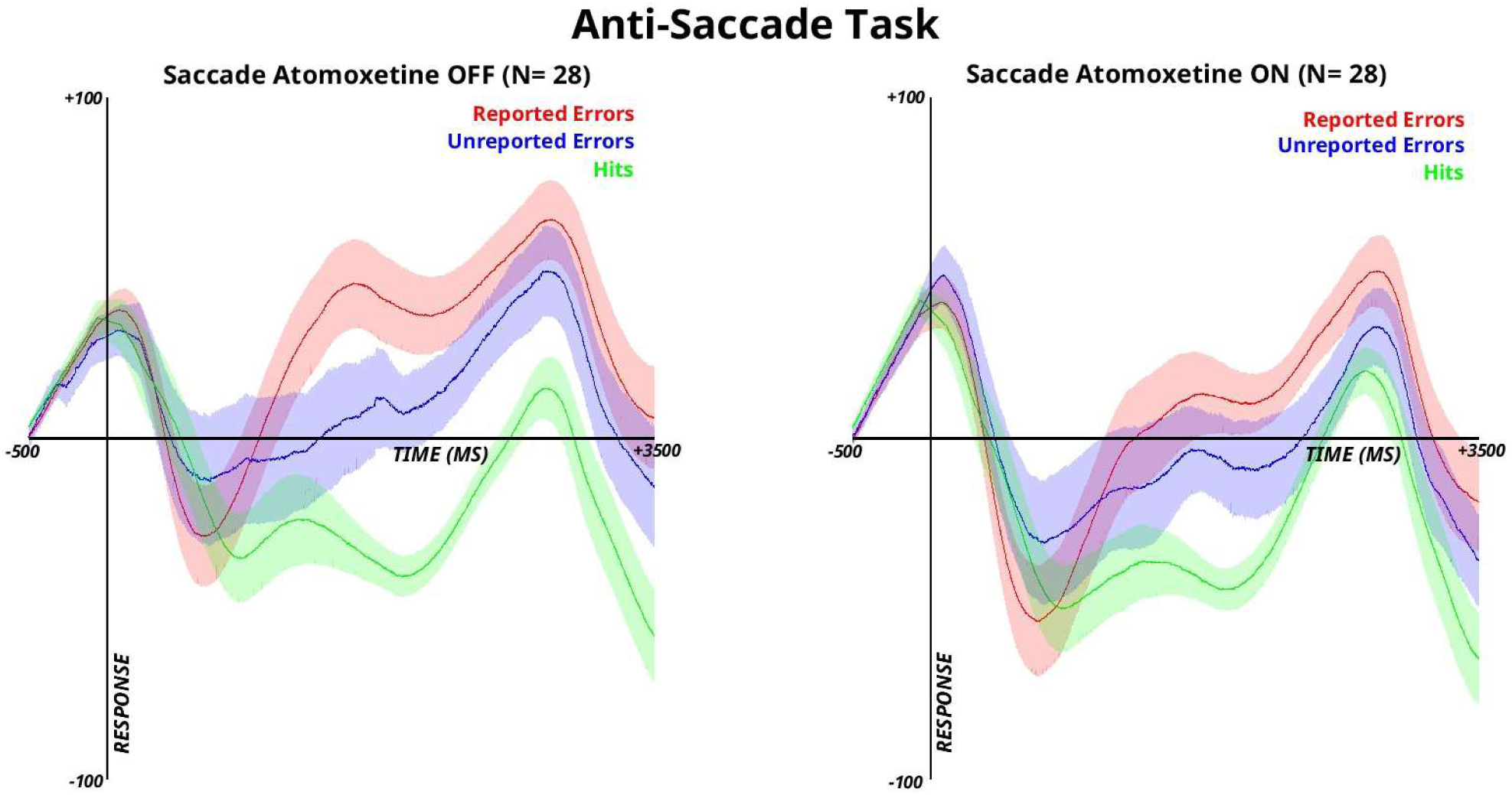
**Grand mean error-related pupil data during the anti-saccade task (n = 28)**. Left: on placebo, we observed phasic increases in autonomic activity for both consciously perceived (red line) and unperceived (blue line) action errors, with a larger increase for reported errors. Right: with atomoxetine, there is a significant decrease in the pupil response for consciously perceived or unperceived errors. Thus, atomoxetine is effectively modulating the phasic autonomic response to action errors.

Planned comparisons then tested for drug effects between sessions. Reported errors were followed by significantly reduced pupil dilation responses under atomoxetine (*M* = 14.73, *SD* = 54.68) compared to placebo (*M* = 39.76, *SD* = 58.70), (*t*(27) = 2.27, *p* = 0.032, *d* = 0.428). Pupil responses to unreported errors did not differ between atomoxetine (*M* = -2.26, *SD* = 62.42) and placebo (*M* = 15.66, *SD* = 65.67), (*t*(27) = 1.28, *p* = 0.210, *d* = 0.242). Neither did the pupil response to correctly identified correct responses (placebo: *M* = -21.94, *SD* = 35.82 vs. atomoxetine: *M* = -24.90, *SD* = 33.21; *t*(27) = 0.45, *p* =0.658, *d* = 0.085).

Together, these results replicate previous reports of greater phasic pupil dilation after action errors, greater phasic pupil dilation after reported vs. unreported action errors, and a direct relationship between the subjective confidence of error detection and the magnitude of the pupil response. Moreover, atomoxetine reduced pupil dilation to consciously perceived errors.

### Behavioral results: anti-saccade task

#### Primary task performance: overall error rate and mean Reaction Time (RT)

Overall error rates did not differ significantly between sessions (placebo: *M* = 11.64, *SD* = 7.15 vs. atomoxetine: *M* = 13.48, *SD* = 9.03; *t*(27) = -1.56, *p* = 0.13, *d* = 0.295). A repeated ANOVA on primary saccadic RTs only showed the main effect of TRIAL ACCURACY (*F*(1, 27) = 47.10, *p* < .001, *η_p_ ^2^* = 0.636), with faster RTs on error than correct trials. This effect was present in both placebo (error: *M* = 268.41, *SD* = 48.97 vs. correct: *M* = 326.96, *SD* = 53.61; *t*(27) = -6.58, *p* < .001, *d* = 1.243) and atomoxetine sessions (error: *M* = 268.85, *SD* = 70.72 vs. correct: *M* = 322.63, *SD* = 56.36; *t*(27) = -5.02, *p* < .001, *d* = 0.949). There was no main effect of SESSION (*F*(1, 27) = 0.07, *p* = 0.800, *η^2^_p_* = 0.002) and no SESSION x TRIAL ACCURACY interaction (*F*(1, 27) = 0.19, *p* = .667, *η_p_^2^* = 0.007).

**Figure 5.**
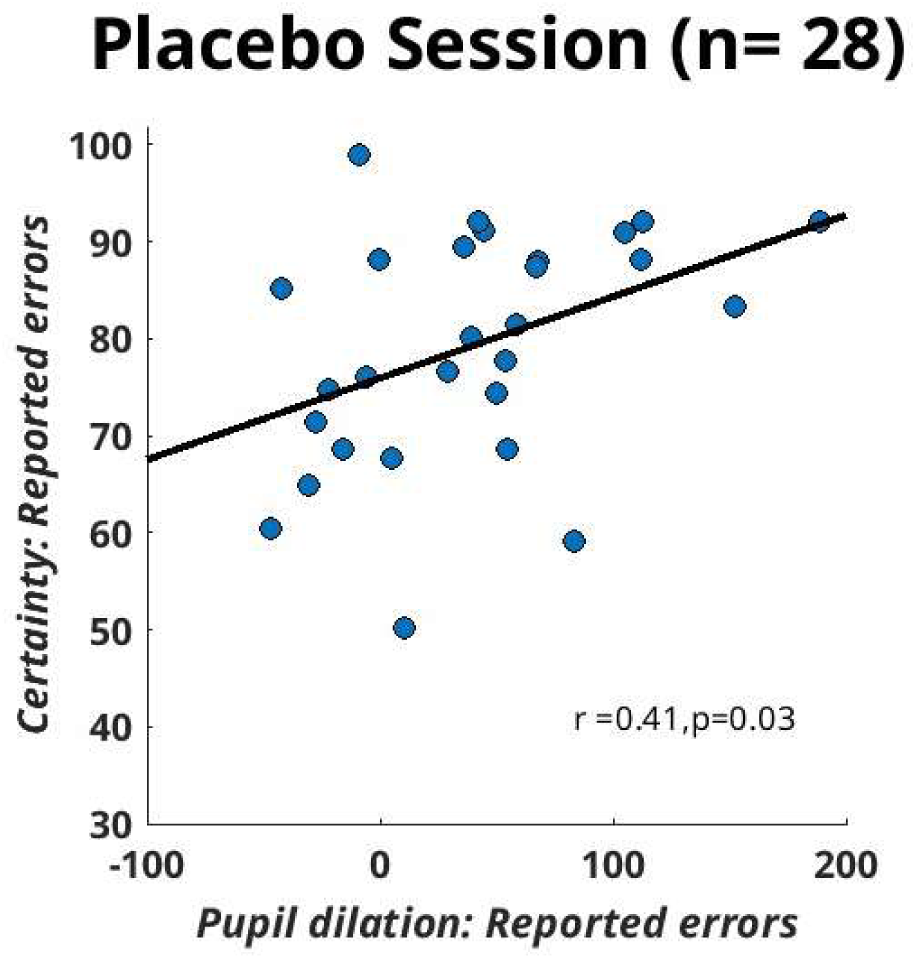
**Correlation between pupil dilation and certainty rating on reported errors.**

#### Error detection rates, rating certainty, and post-error slowing

Performance-monitoring metrics based on error detection queries are shown in **Table 2**. Error detection rates did not differ between sessions (placebo: *M* = 76.30, *SD* = 13.98 vs. atomoxetine: *M* = 76.60, *SD* = 11.38; *t*(27) = -0.15, *p* = 0.884, *d* = 0.028). False alarm rates were also comparable (placebo: *M* = 14.73, *SD* = 16.01 vs. atomoxetine: *M* = 15.50, *SD* = 14.35; *t*(27) = -0.33, *p* = 0.742, *d* = 0.063).

A two-way repeated measures ANOVA on rating certainty revealed a main effect of TRIAL TYPE (*F*(1.69, 45.59) = 14.24, *p* < .001, *η_p_^2^* = 0.345), with highest certainty for reported errors (*M* = 79.08), lowest for unreported errors (*M* = 67.38), and intermediate values for hits (*M* = 72.25). There was no main effect of SESSION (*F*(1, 27) = 0.00, *p* = 0.964, *ηp^2^* < 0.001) and no SESSION x TRIAL TYPE interaction (*F*(1.46, 39.33) = 0.437, *p* = 0.586, *η_p_^2^ =* 0.016). Planned comparisons confirmed no session differences for hits (placebo: *M* = 72.94, *SD* = 16.46 vs. atomoxetine: *M* = 71.57, *SD* = 16.01; *t*(27) = 1.01, *p* = 0.322, *d* = 0.191), reported errors (placebo: *M* = 79.24, *SD* = 12.04 vs. atomoxetine: *M* = 78.91, *SD* = 14.20; *t*(27) = 0.22, *p* = 0.828, *d* = 0.042) and unreported errors (placebo: *M* = 66.62, *SD* = 19.91 vs. atomoxetine: *M* = 68.14, *SD* = 16.16; *t*(27) = -0.46, *p* = 0.652, *d* = 0.086).

PES results are depicted in **Table 2**. A two-way repeated measures ANOVA on PES revealed a main effect of ERROR TYPE (*F*(1, 27) = 6.97, *p* = .014, η_p_*^2^* = 0.205) with greater PES following reported (*M* = 12.16) unreported errors (*M* = 3.31). In the placebo session, PES significantly greater after reported (*M* = 10.33, *SD* = 12.30) than unreported errors (*M* = 2.45, *SD* = 12.85), (*t*(27) = 2.93, *p* = 0.007, *d* = 0.554), replicating our prior findings (Wessel et al., 2018). In the atomoxetine session, the same pattern was numerically present (reported: *M* = 14.00, *SD* = 29.37 vs. unreported: *M* = 4.16, *SD* = 13.88) but not significant (*t*(27) = 1.57, *p* = 0.127, *d* = 0.297). There was no main effect of SESSION (*F*(1, 27) = 0.57, *p* = 0.46, *ηp^2^* = 0.021), and no interaction (*F*(1, 27) = 0.08, *p* = 0.78, *ηp^2^* = 0.003).

Taken together, atomoxetine did not significantly affect primary task performance, error detection rates, rating certainty, or PES (See **Table 1**-**2**). The established pattern for greater PES following reported versus unreported errors, which we here replicated in the placebo session, was absent on atomoxetine.

**Table 1.** Primary task performance for overall error rates (%) and mean RT (ms). Statistics reported are mean values (± SD).

|  | Primary Task Performance |
| --- | --- |

| **Session** | *Overall error rate* | *Correct RT* | *Error RT* |
| --- | --- | --- | --- |
| **Placebo** | 11.64 (7.15) | 326.96 (53.61) | 268.41 (48.97) |
| **Atomoxetine** | 13.48 (9.03) | 322.63 (56.36) | 268.85 (70.72) |

**Table 2.**
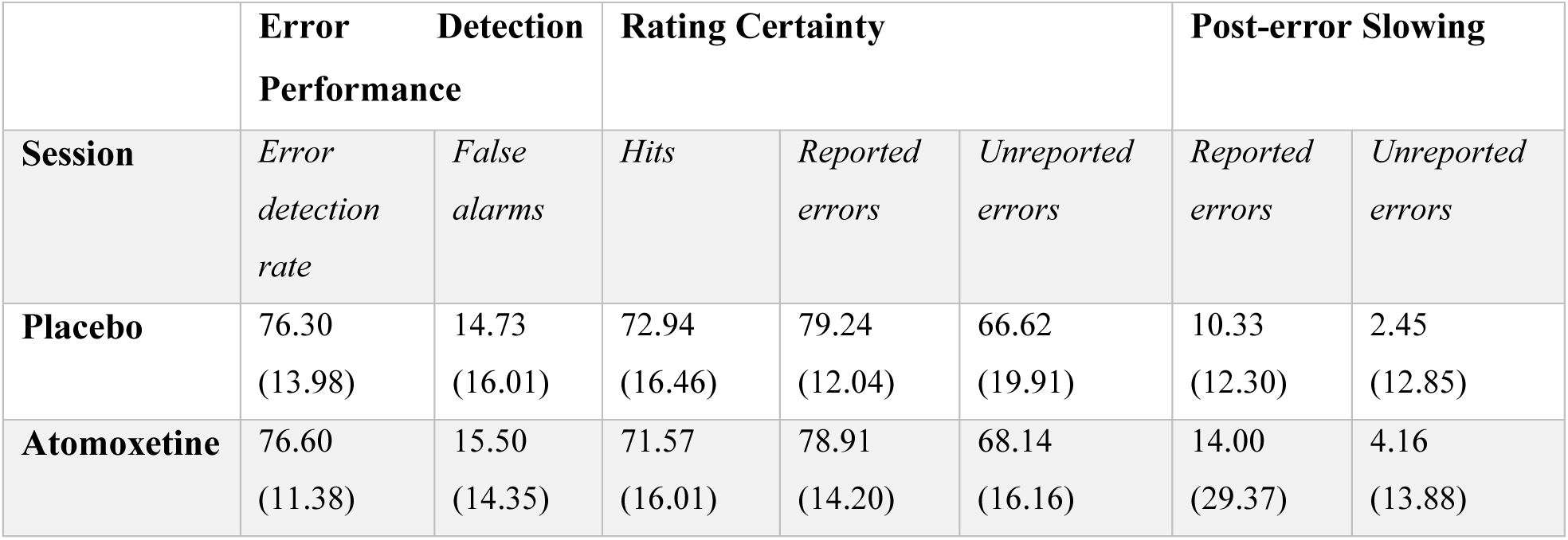
Task performance based on trials included the error detection query. Statistics reported are mean values (± SD).

## Discussion

Prior studies have illustrated that the phasic autonomic response and conscious error detection are tightly linked (Wessel et al., 2011; Ullsperger et al., 2010; Shalgi et al., 2009; Hajcak et al., 2003). The causal nature of this relationship was hitherto unexplored. In the present study, healthy older adults performed an anti-saccade task while we administered the sNRI atomoxetine (80 mg) in a double-blind, placebo-controlled fashion, with the goal of modulating the phasic autonomic response to errors.

In line with our previous work, consciously perceived action errors were associated with a larger phasic autonomic response, and the degree of this autonomic error response was correlated with the subjective certainty of conscious error detection (Figures 4 **and 5**). Our main hypothesis pertained to the differences in error detection behavior. If the autonomic system contributes evidence to the conscious error detection process, we would expect conscious error detection to be altered under atomoxetine, alongside the observed alterations to the phasic autonomic response. However, if the phasic autonomic arousal was a consequence of conscious error detection, we would expect no differences to conscious error detection performance, despite the differences in autonomic response induced by atomoxetine.

Our results clearly favor the second interpretation (**Table 2**). There were no differences in the rate of conscious action error detection, the false alarm rate, or the overall accuracy of action error detection. Moreover, there was no difference in the subjective rating certainty on any of the trial types. Primary task performance was also unaffected (**Table 1**), as neither error rate nor saccadic reaction time to any of the trial types differed between the two sessions. Post-error slowing was observed to equal degrees in both sessions, with greater post-error slowing on consciously perceived errors (though this effect was absent on atomoxetine). Taken together, these findings suggest that the observed phasic increase in autonomic arousal after errors is a consequence, and not a cause, of conscious error detection.

Atomoxetine exhibits population-specific behavioral effects: while it reliably enhances performance in younger individuals and children diagnosed with ADHD (Chamberlain et al., 2007; Wu et al., 2021; Shang & Gau, 2012; Barkley et al., 2007), its benefits appear minimal in healthy older adults. To our knowledge, this study provides the first test of atomoxetine’s effects on task performance in this age group. Our results suggest that atomoxetine’s cognitive benefits are largely condition-dependent and may not extend to healthy adults.

To summarize, we measured the effects of the sNRI atomoxetine on phasic changes in pupil diameter and performance monitoring in a group of healthy older adults, to assess the direction of causality between the phasic autonomic response and error awareness. While we observed an effect of atomoxetine on the phasic autonomic response, we did not find a significant effect for error awareness. Thus, our results suggest that conscious error detection causes the autonomic response to errors, not vice versa.

## Acknowledgements

This research was supported by a grant from the Aging Mind and Brain Initiative to JRW. TNJ was supported by the Andrew H. Woods Professorship at the University of Iowa and JRW was supported by the Clement T. and Sylvia H. Hanson Family Chair at the University of Iowa.

